# Purification and characterization of a novel anti-tumour galactose binding lectin from *Euphorbia caducifolia* latex

**DOI:** 10.1101/2023.03.10.532027

**Authors:** Kusuama Venumadhav, Kottapalli Seshagirirao

## Abstract

A galactose specific lectin was isolated from the latex of *Euphorbia caducifolia* to investigate the therapeutical aspects protein. Protein was purified from latex serum with two different affinity matrices and confirmed to contain 8 isoforms called isolectin with a molecular weight of 32 and 32.5 kDa. Preliminary studies state the specificity towards galactose, but its affinity with lactose is more favourable even at low concentrations. Circular spectrometry suggests the predominance β conformation is high in comparison to others as like most of the other latex lectins. Haemagglutination activity of the protein with different thermal and pH ranges shows maximum activity at pH 6-8 and 60-70°C. In therapeutically, it is noticed that the galactose specific lectin showed mitogenicity with human macrophages and anti-cancer properties on human cervical cancer cells.

## 1. Introduction

Lectins have been used to investigate the immunology of Red Blood Cells. Recently, Anti tumor and antiangiogenic activities were noticed with several glycoproteins from Euphorbia species [1]. Now, it is important to explore the new moieties to treat the tumour cells apart from chemical as well as radiation therapies. Many studies have shown the importance of small peptides and more protein in treating tumour activities.

Seed lectins from Euphorbiaceae were studied by Stillmark (1888) initiates the immunology of lectins [2]. Further elevates the lectin studies on isolation, purification, and their biological applications. Lectins have been detected and isolated from different species of all living sources but the majority from angiosperms. *Azolla caroliana*, a pterridophyte has been reported for the presence of a lectin [3]. Seeds of angiosperms are the major contributors of the lectins though several lectins have also been detected and isolated from various parts of the plants. For example, from leaves of *Aloe arborescens* [4], *Dolichos biflorus* [5], *Griffonia simplicifolia* [6]; Barks of *Robinia pseudoacacia* [7], *Sambucus nigra* [8]; Sieve saps of *Castanea sativa, Salix alba* [9]; Bulbs of *Galanthus nivalis* [10], *Leucojum aestivum* [11], *Narcissus spp*. [12], *Tulipa gesneriana* [13]; Tubers of *Bryonia dioica* [14], *Trichosanthes kirilowii* [15].

Euphorbia caducifolia, a semi-arid plant native to the Indian subcontinent and grows in rocky areas, and inhabitant used the plant for various therapeutical purposes [16]. Latex biochemistry studies of the plant suggest the presence of galactose binding lectins [17]. In an interest to the later report, this study aimed to explore the biological importance of the galactose specific lectin along with its biochemical characterization. Moreover, evaluated the different matrices in the purification of lectin.

## 2. Materials and Methods

### 2.1 Purification of Galactose-binding Protein

The latex of *Euphorbia caducifolia* was collected and frozen in liquid nitrogen; Frozen serum was thawed and centrifuged at 20,000 x g for 1 hour at 4°C. The liquid portion was separated and repeated the freezing, thawing and centrifugation process to get clear serum. The separated clear serum was used for experiments or stored below -20°C until required. The crude serum proteins were precipitated with cold acetone (overnight at -20°C) and centrifuged at 10,000 x g for 10 minutes. The protein pellet was dissolved in phosphate buffer saline and used for experiments or stored at -20°C.

### 2.2 Preparation of Affinity matrix

The Affinity matrices were prepared according to Seshagirirao method [18]. In brief, 100 g of seeds of (Guar gum and Leucaena leucocephala seed gum) was blended at low speed for few seconds. The seed hulls were removed, and kernels (60g) were milled into fine powder. The powder was further sieved through 1mm mesh. Next, the powder was mixed with an emulsion of Epichlorohydrin (99%) in 3 M NaOH. The mixture was incubated for 24 h at 40°C and later for 6 h at 70°C. The formed matrices were soaked and washed several times with distilled water. Finally, the recovered gel was transferred to 10 mM phosphate buffer saline, pH 7.2 containing 0.02% of sodium azide. Obtained protein was estimated by Lowry’s method

### 2.3 Purification of Galactose-Binding Protein

The purification of galactose binding protein by affinity chromatography was performed at 4°C. 2 mL of latex serum (50μg) was loaded onto matrix column (3 × 7 cm, ca 50 mL), previously equilibrated with phosphate buffer saline pH 7.2 with a flow rate 15 mL/h. The column was washed with it until the effluent gets < 0.02 absorbance at 280 nm. The protein adsorbed was eluted with 0.2M lactose in Phosphate buffer saline. The purity of high protein fractions was verified by SDS-PAGE, pooled and dialyzed against phosphate buffer saline and further determined the isoelectric nature of the protein by 2D electrophoresis.

### 2.4 Carbohydrate Estimation and monosaccharide composition

Total carbohydrate was estimated using Dubois method [19]. The glycosylated carbohydrates to the protein were studied using GC-MS analysis. Lectin (50 μg) was hydrolyzed with 4 M TFA by incubating for 6 h at 80°C. Hydrolysate was dried with methanol under speed vac. The residue was dissolved in pyridine and derivatized with silylating reagent (BSTFA and 1% TMCS) in 1:2 ratio at 40°C for 2h. Monosaccharide derivatives were extracted with dichloromethane and analyzed for composition using GC-MS [20]. Quatitized the monosaccharides using a set of standards.

### 2.5 Haemagglutinating activity

The haemagglutinating activity of the galactose binding protein was determined with minor modifications of [21] method. 100 μl of the protein sample (final conc. 1μg) was serially diluted in a microtitre plate, and 100 μl of trypsinized 4% erythrocytes were added to each well. The agglutination was observed visually after incubation of the plate for one hour at 37°C. The highest dilution which showed positive haemagglutination was considered as the Titre. The amount of protein presents in this dilution represents the minimum quantity of protein required for agglutination and is defined as one unit. The specific activity is the number of units per mg of protein.

### 2.6 Sugar Inhibition Assay

50 μl (0.4 M) of serially diluted carbohydrate solution was mixed with 50 μl protein containing 8 haemagglutination units in a microtitre plate and incubated at room temperature for 30 min. 100 μl of 4% trypsinized human O group erythrocyte suspension was added to the incubated solution for 1 h at 37°C. The inhibition concentration of the sugar was recorded as the minimum concentration of sugar required for complete inhibition of 2 haemagglutinating units with 2% of the erythrocytes.

### 2.7 Temperature and pH stability

The lectin (50 μg) was incubated at different temperatures (4°C, 30°C, 40°C, 50°C, 60°C, 70°C, 80°C and 90°C) for 10 minutes and was brought to room temperature. The ability of haemagglutination was tested to determine the temperature. The lectin (50 μg) was incubated at different pH (5, 6, 7, 8, 9 and 10) for 30 minutes and the haemagglutination activity was tested to determine optimum pH.

### 2.8 CD spectroscopy studies

Circular dichroism spectra were recorded to examine the secondary confirmations of the lectin using JASCO spectrometer. The lectin was placed in the 2 mm Quartz cuvette with a concentration of 2 HU of protein and recorded spectra. Spectra measured for Far UV region (250-190 nm) and Near UV region (300-250 nm) with 50 nm per minute and accumulated 3 scans. The secondary structure was analyzed using Dichroweb online software. Also, the effect of the temperature and pH on the unfolding of the lectin was also studied.

### 2.9 Partial peptide sequence

Partial peptide sequence of the lectin was obtained from the in-gel digestion and MALDI MSMS. The peptide sequence was performed by “Sandor life sciences pvt ltd” and obtained peptides were identified by peptide mass fingerprinting and MSMS MASCOT software.

### 2.10 MTT assay

RAW 264.7 was used for the experiment. The 1×10^4^ cells were inoculated and incubated for 24 h in the CO_2_ incubator. The lectin was added to the cells at different concentrations and incubated for various time points. After incubation, MTT, 5 mg/mL was added and incubated in the dark for 4 h. Crystallized formazan was extracted with DMSO and estimated at 590nm of Absorbance. Cell proliferation was calculated with control cells (without lectin) [22].

### 2.11 Antitumour activity and Cell Migration Assay

Mouse monocyte macrophages (J774a.1), Human cervical cancer cells (HeLa). The cells (10,000) were inoculated in 96 well plates and incubated for 24 h in the CO_2_ incubator. Then, the cells were treated with different concentrations protein and incubated for various time points. After time points, 3(4,5-dimethyl thiazol-2-yl)-2,5-diphenyl tetrazolium bromide (MTT) was added as 5 mg/mL and incubated further for 4 h. 100 ul of DMSO was added to the wells and absorbance taken at 590nm.

Monolayer cells of HeLa cells were grown to confluence in a 12 well plate. A scratch was created by using PCR micro tip in the middle of the monolayer well. Detached cells were removed by washing with incomplete media twice. Then the cells were treated with EC lectin with 50, 100 and 200 μg /ml and untreated was considered as control. The plate was incubated, and pictures of the gap area were taken at different time points to observe the migration of the cells towards the gap.

## 3. Results and Discussion

### 3.1 Purification of Galactose Binding Lectin

Lectins are glycoproteins with carbohydrate moieties and can agglutinate the blood. Lectins show specificity towards various sugars. Galactose-binding lectins are peculiar to the galactose and their derivatives as the name says. Galactose-binding lectins can be purified by affinity chromatography using galactose-specific matrices. Seshagirirao and Prasad (2001) reported the presence of galactose-specific lectins in Euphorbia caducifolia and other Euphorbia species. Lectins from *Euphorbia caducifolia* latex was isolated by applying on galactose affinity matrices. Affinity matrices were prepared and evaluated with snake gourd lectin. Applied crude serum was eluted with 0.2 M Lactose and collected the protein (Fig. 1). Two matrices were used to assess the yield of the protein. Purified protein with a yield of 60% was observed with CLGG matrix, and 82% of yield was noted with CLLSG (Table 1). CLLSG shows more affinity and gives high purification folds compared to the other matrices like Guar gum, cashew nut and egg matrices [18] and also with Sepharose 4B.

**Table 1.** The yield of galactose-binding protein from *Euphorbia caducifolia* latex on different affinity matrices.

| Purification step | Protein (mg) | Specific Activity | Total Activity (titre X mg) | Yield (%) |
| --- | --- | --- | --- | --- |
| Crude Protein | 50 | 2560 | 128000 | 100 |
| CLGG | 4.5 | 17066 | 76797 | 60 |
| CLLSG | 6.2 | 17066 | 105809.2 | 82 |

**Fig. 1.**
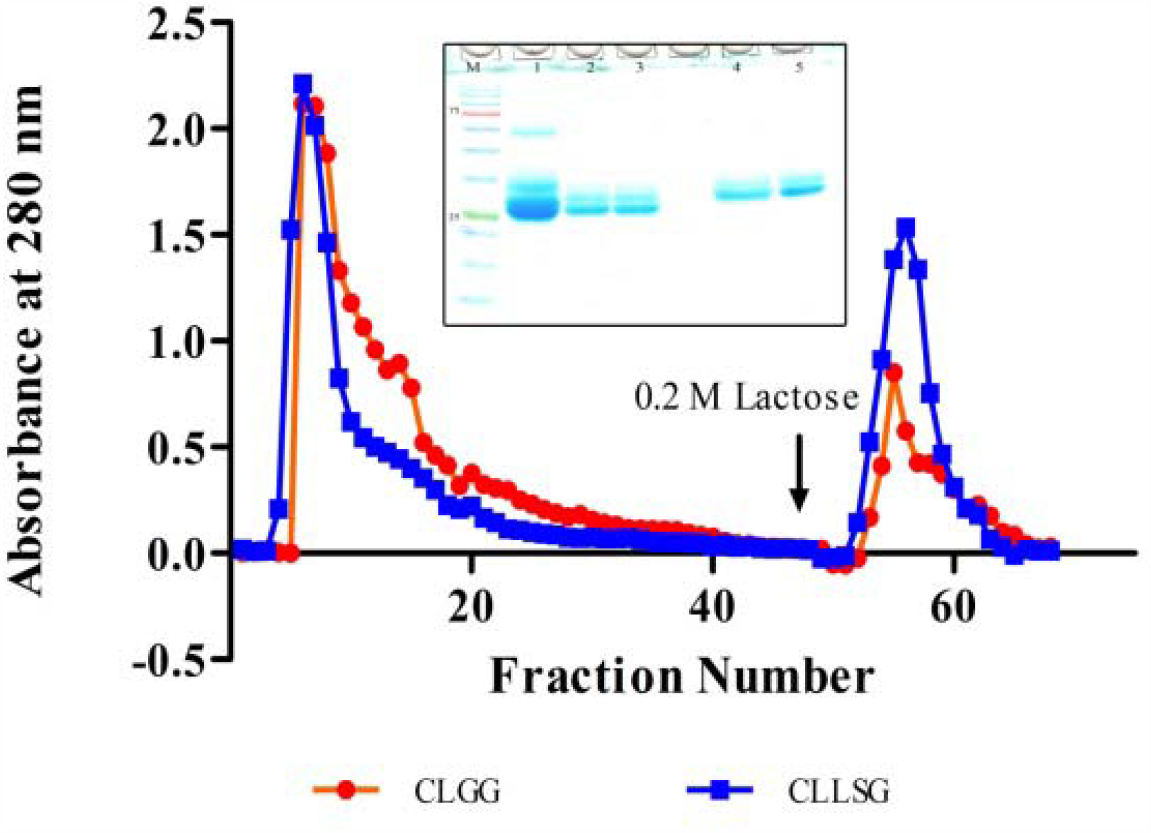
Affinity Chromatography of *Euphorbia caducifolia* Latex serum on CLGG, CLLSG column (3×7 cms) at 4 ^0^C.; Insert SDS-PAGE of Purified Galactose-binding protein from *Euphorbia caducifolia* latex : M- Protein marker, 1- Crude protein, Purified protein 2-CLGG, 3- CLLSG, and purified protein with reducing agent β-Mercaptol, 4- CLGG, 5- CLLSG.

Previously, it has been reported the isolation of ECL using Sepharose 4B column with a yield of (67%) which is more as observed with CLGG and less with CLLSG. It shows that the use of CLLSG for isolation or purification of galactose binding lectin is more efficient than other columns. Further purified protein was run over SDS-PAGE to check the lectin purity (Fig. 1 (inset)) and lyophilized until the studies its biochemical properties. From the electrophoresis analysis, it has been observed the protein fraction consist of 8 isolectins molecular weight varying with 32 kDa (IL1, pI 4.3 & IL-2, pI4.6) and 32.5 kDa (6 forms) and shows the charge heterogeneity in the range of pI 4.3-5.8.

### 3.2 Haemagglutinating activity and carbohydrate specificity

The haemagglutination activity of ECL was observed more with O compared to other blood groups (Table 2). ECL did not show much difference in agglutinating trypsinized red blood cells of Rh positive A, B, and AB groups but agglutinated almost 6 folds with Rh O group. It indicates the specificity of the ECL towards the O type as other Euphorbia spp. [23].

**Table 2.** Haemagglutination of Lectin with trypsinized human Rh^+^ Erythrocytes.

| Erythrocyte | Specific Activity (U/mg) |  |
| --- | --- | --- |
|  | Crude Protein | Purified Protein |
| A<sup>+</sup> | 400 | 2560 |
| B<sup>+</sup> | 400 | 2560 |
| AB | 400 | 2560 |
| O<sup>+</sup> | 2560 | 17066 |

All lectin are carbohydrate specific and inhibition the agglutination of RBC in the presence of their specific sugar where binding sites of the lectin blocked. ECL was observed specificity with different carbohydrates (Table 3), galactose and galactose derivatives (Table 4). We noticed the delay in agglutination with melibiose and raffinose where the galactose in 1→6 glycosidic linkage but observed no agglutination with lactose where the glycosidic linkage is involved is the 1→4 type. Apart from the specificity of the lectin to the sugar, and also express its specificity with linkages of specific monomer in oligosaccharides. O-Nitrophenyl galactoside also inhibits the agglutination as galactose majorly binds with lectin sites. Reports say that most of the latex lectin from Euphorbia are specific in galactose moiety either direct or its derivatives [24,25].

**Table 3.** Sugar Inhibition Assay with different sugars and oligosaccharides.

| S.No | Carbohydrate | Agglutination Activity | Galactose present (Yes/No) | Saccharide |
| --- | --- | --- | --- | --- |
| 1 | Agar | Present | Yes | Polymer |
| 2 | Agarose | Present | Yes | Polymer |
| 3 | Arabinose | Present | No | Mono |
| 4 | Dulcitol | Present | Yes | Mono |
| 5 | Fructose | Present | No | Mono |
| 6 | Fucose | Present | Deoxy Galactose | Mono |
| 7 | Galactosamine | Absent | Yes | Mono |
| 8 | Galactose | Absent | Yes | Mono |
| 9 | Glucose | Present | No | Mono |
| 10 | Inositol | Present | No | Mono |
| 11 | Mannose | Present | No | Mono |
| 12 | N-Acetyl galactosamine | Absent | Yes | Mono |
| 13 | N-Acetyl glucosamine | Present | No | Mono |
| 14 | Rhamnose | Present | No | Mono |
| 15 | Salicin | Present | No | Mono |
| 16 | Xylose | Present | No | Mono |
| 17 | Lactose | Absent | Yes | Di |
| 18 | Melibiose | Present (Slow) | Yes | Di |
| 19 | Raffinose | Present (Slow) | Yes | Tri |
| 20 | O-Nitrophenyl galactopyranoside | Absent | Yes | Derv. |

**Table 4.**
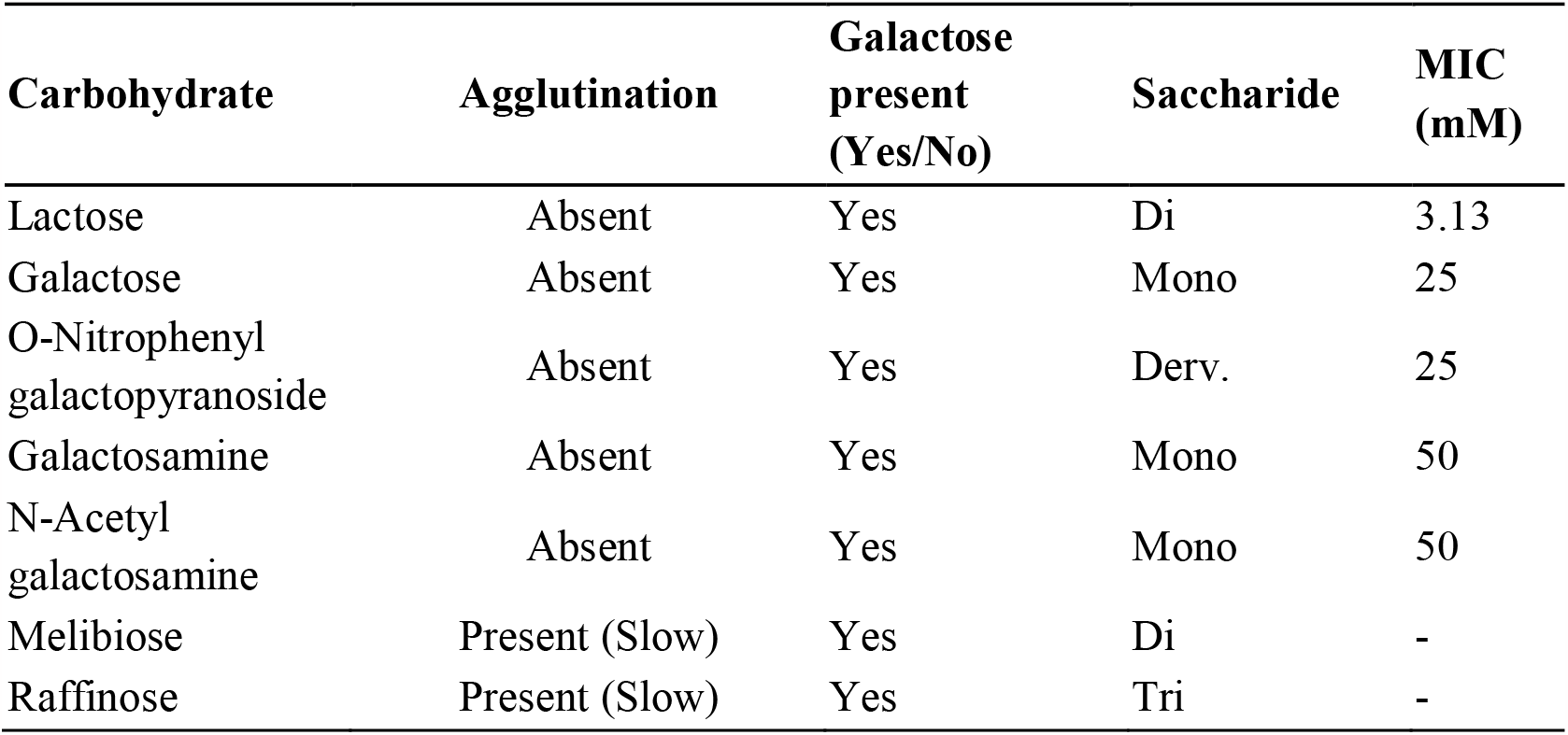
Sugar Inhibition Assay with different Galactose derivatives and their MIC.

| **Carbohydrate** | **Agglutination** | **Galactose**<br>**present**<br>**(Yes/No)** | **Saccharide** | **MIC**<br>**(mM)** |
| --- | --- | --- | --- | --- |
| Lactose | Absent | Yes | Di | 3.13 |
| Galactose | Absent | Yes | Mono | 25 |
| O-Nitrophenyl<br>galactopyranoside | Absent | Yes | Derv. | 25 |
| Galactosamine | Absent | Yes | Mono | 50 |
| N-Acetyl<br>galactosamine | Absent | Yes | Mono | 50 |
| Melibiose | Present (Slow) | Yes | Di | - |
| Raffinose | Present (Slow) | Yes | Tri | - |

### 3.3 pH and temperature effect

Lectin isolated was observed for the influence of pH and temperature on activity (Fig. 2). ECL shows a broad range of activity in pH as well as temperature. Lectin activity notified at pH range 6.0-9.0 but showed maximum activity at pH 8.0. Maximum activity or association constants of the several lectins were found at neutral or slightly alkaline pH [26]. Like most of the Euphorbiacae latex lectins, ECL also inactive above pH 10.0. Temperature plays a good role on activity as well as the stability of a protein apart from pH as physical conditions. *E. caducifolia* lectin is heat stable and shown maximum activity at 60°C and 70°C, and gradually activity ceased above 90°C.

**Fig. 2.**
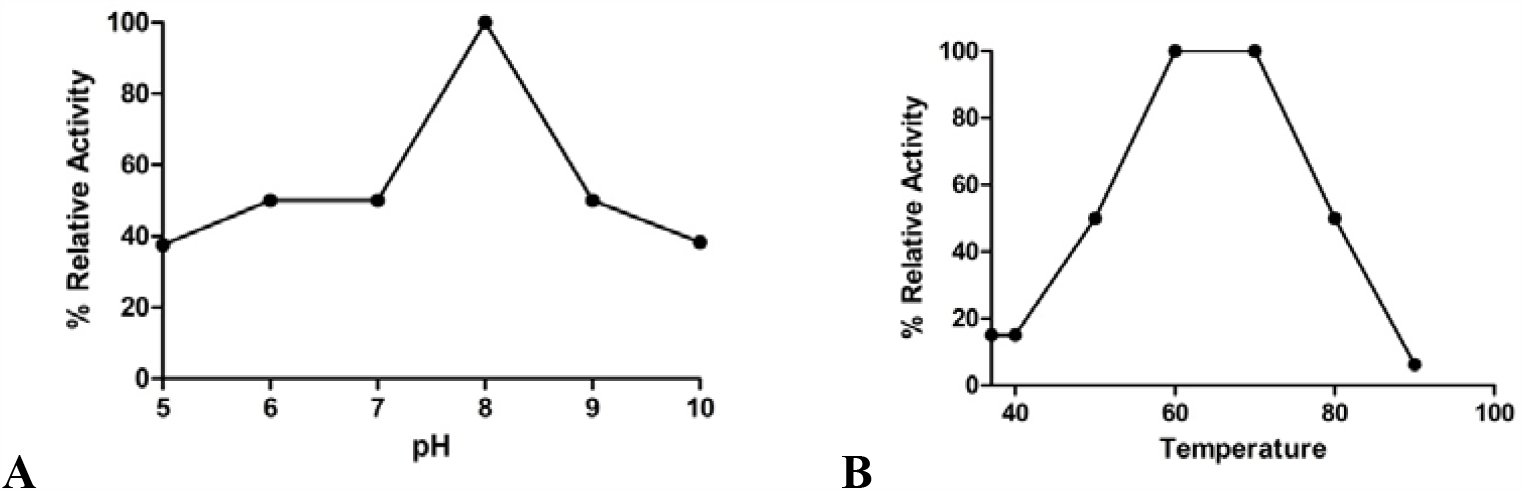
Physicochemical parameters influence on Lectin haemagglutinating activity (A) pH, (B) Temperature

### 3.4 Glycoprotein characterization

Glycoprotein character of the lectin was identified by the Schiff periodic base on PAGE. The carbohydrate portion of the lectin was estimated to be 11.82%, and this was almost nearer to the previous reports. Carbohydrate content of the lectin was very similar to the amount present in other species of the genus. Further studies revealed the presence of monosaccharides covalently bound to the protein (Table 5). Sugars found in the protein allude to structural conformations of protein as well as its biological importance.

**Table 5.** Sugar composition in the Lectin

| Sugar | Residue/Molecule |
| --- | --- |
| Fuc | 11.3 |
| Gal | 8.3 |
| Glu | 0.6 |
| Man | 8.9 |
| GLcNac | 12.7 |
| Xyl | 1.4 |
| Total sugar (%) | 11.4 |

### 3.5 CD Spectroscopy Studies

Structural confirmation of the lectin was studied by Circular Dichroism spectrometry. Spectra of the native lectin in the near UV and far UV was recorded at 25°C. Far UV spectrum observed with 2 positive peaks at 195 and 230 nm, and 2 negative peaks at 210 and 218 nm. Secondary structure information of the lectin was analyzed by software available at DICHROMEWEB. The secondary structural content obtained are 1% of regular α helix and 6% of disordered α helix (Total α helix of 7%), 21% of regular β sheet and 14% of disordered β sheet (Total β sheet of 35%), 24% β turns and 34% of unordered structures (Fig. 3). Overall CD spectrum of the lectin indicates that the predominant of β sheet protein to minor α helical structure of the protein. Influence of the temperature and pH on the secondary structure of ECL also studied using CD spectroscopy. As affecting the RBC agglutination, temperature, as well as pH, alters the structure of the lectin.

**Fig. 3.**
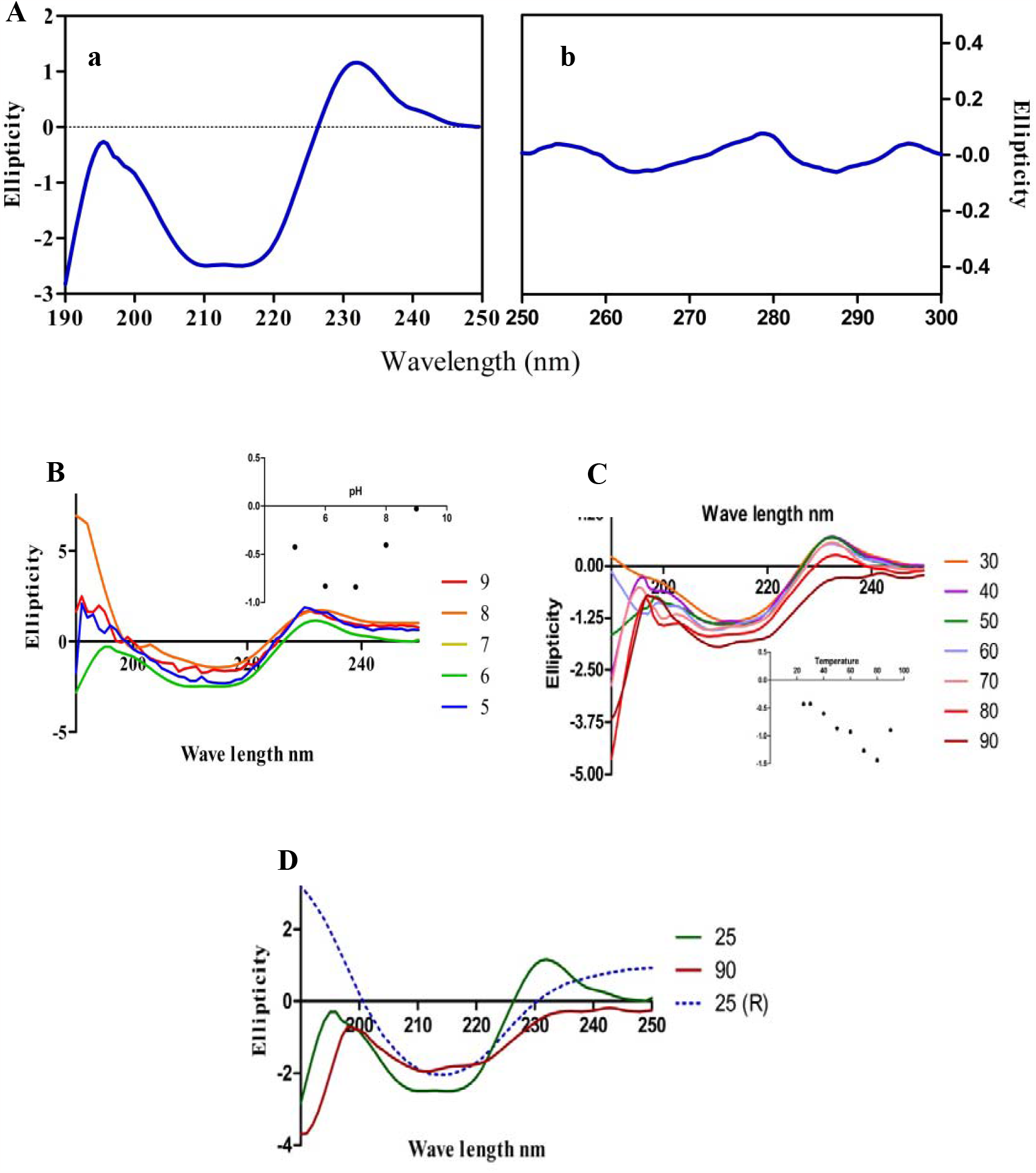
A) Secondary confirmation of the lectin by CD spectrometry, a) Far UV, b) Near UV; Influence of B) pH and C) Temperature on structure, D) Spectrum of cooled lectin after incubated at 90°C.

### 3.6 Partial peptide sequence

Purified lectin was subjected to peptide sequence to discover the fragments or domains of the protein. Peptides gained many attentions towards medical and biotechnology aspects, and over 7000 peptides have been identified as they play important roles in human physiology [27]. Much more, peptide research on therapeutic experiencing a commercial renaissance. 3 Major and significant peptides were identified from a total of 8 obtained peptides from Euphorbia caducifolia lectin (Fig. 4). 3 peptides were sequenced for amino acids by MS/MS analysis and blasted the sequences for identification through NCBI protein database. Not many reports have been found on Euphorbia latex or seed lectin protein sequence or peptide sequence. Previously, only peptide sequence reported from E. characias and E. marginata lectin [29] and total protein sequence of a mannose-binding lectin from E. tirucalli was reported recently [30]. The peptide sequence of Euphorbia lectins was not identical and didn’t show any conservative domains of the family.

**Fig. 4.**
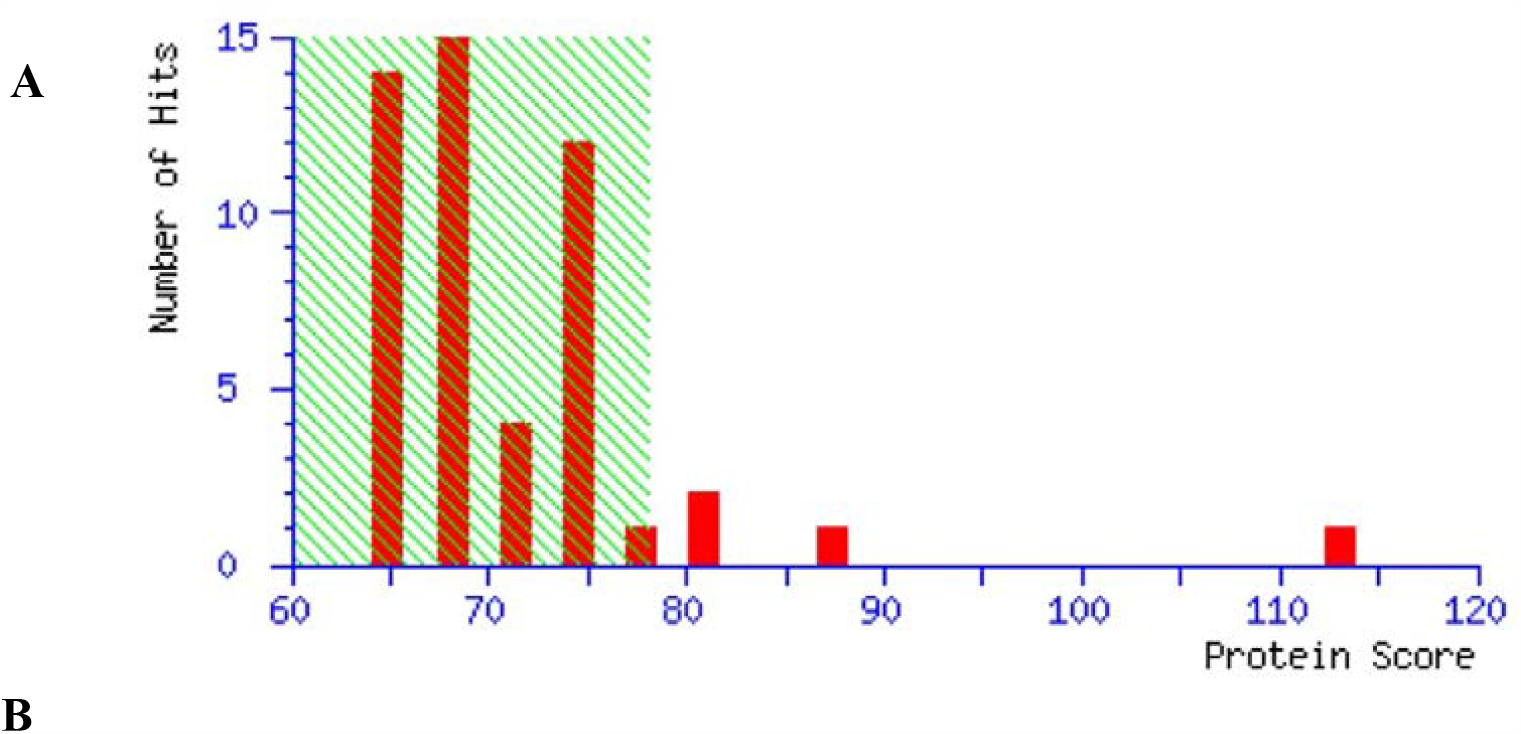
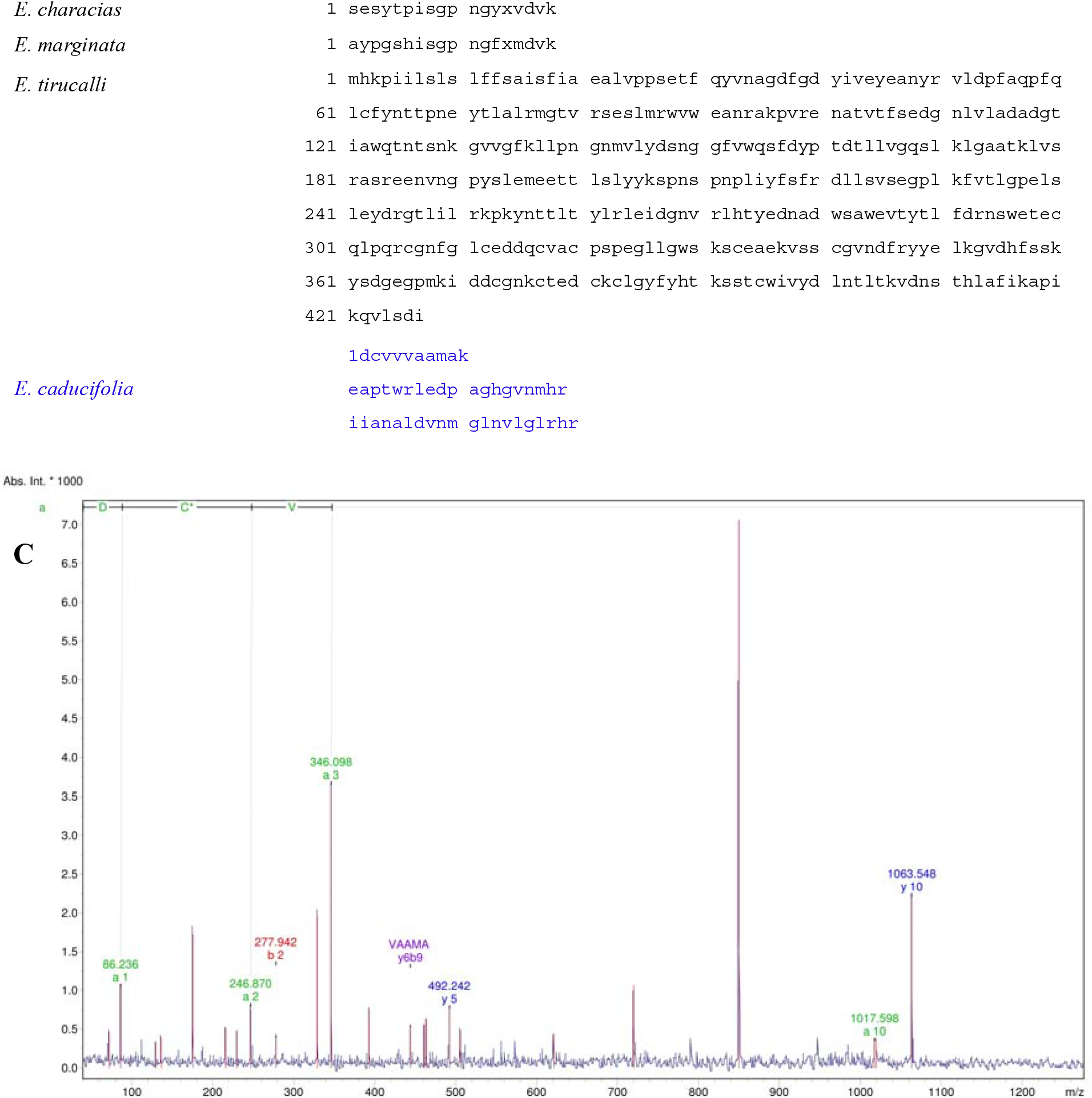
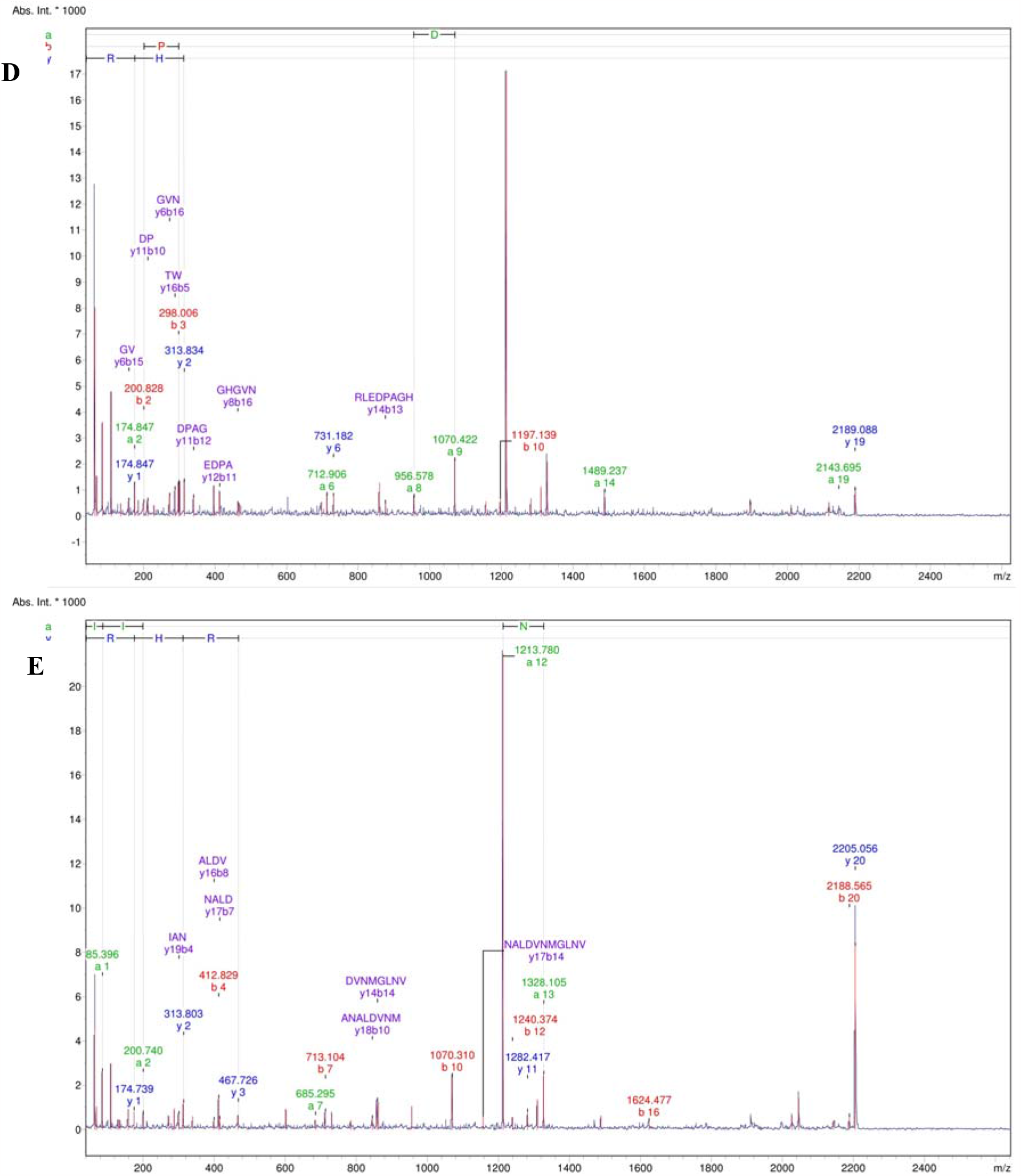
Partial peptide sequencing of Lectin, A) Protein score histogram (78*), B) Euphorbia lectins peptide sequences; Peptide mass spectrum of masses C) 1063, D) 2189, E) 2205. * Protein score greater than 78 are significant (p<0.5)

### 3.6 Hemolytic activity

Aside of agglutination, prolong incubation of RBC with latex serum exhibits the hemolytic activity. *Euphorbia caducifolia* lectin did not show any hemolytic activity with RBC used in haemagglutination assay. Some of the seed lectins like *Sterculia foetida L* and *Clitoria fairchildiana* were reported for their hemolytic activity as most of the lectins in particular latex lectin does not display such activity. The absence of hemolytic activity of the drugs or molecules imparts therapeutic attention as they do not harm the biological system. Lectins are using in various therapeutic application due to non-hemolytic nature as like ECL.

### 3.7 Cell proliferation and Anti-tumour activity

Lectins gained attention because of their proliferative activity on different cell types due to cell surface carbohydrate recognition variabilities [28]. Euphorbia latex lectins can act as mitogens which induce the mitosis of the cells. Latex lectins from *E. nerifiolia* and *E*.*marginata* were reported for their mitogenic activity [29]. In similar *E. caducifolia* also exhibited a mitogenic activity which was studies by cell proliferation. Mitogens induce the mitotic division of the cell which ultimately leads to the cell proliferation. *E. caducifolia* lectin treated cells were showed an increase in the cell number at all time points and all concentrations (Fig. 5). Cell number increased with increasing concentration of the lectin and this might be due to the mitogen activity of lectin. Maximum activity showed at 3 and 24 h of incubation, but it slightly decreased at 100 μg/ml concentration in relative to the other time points after 24 h of treatment. Proliferation ranged in between 0-50% more as compared to the untreated cells (control), and it considered as ECL can act as a Mitogen. The diverse role of lectins in cell proliferation and differentiation is still undelivered.

**Fig. 5.**
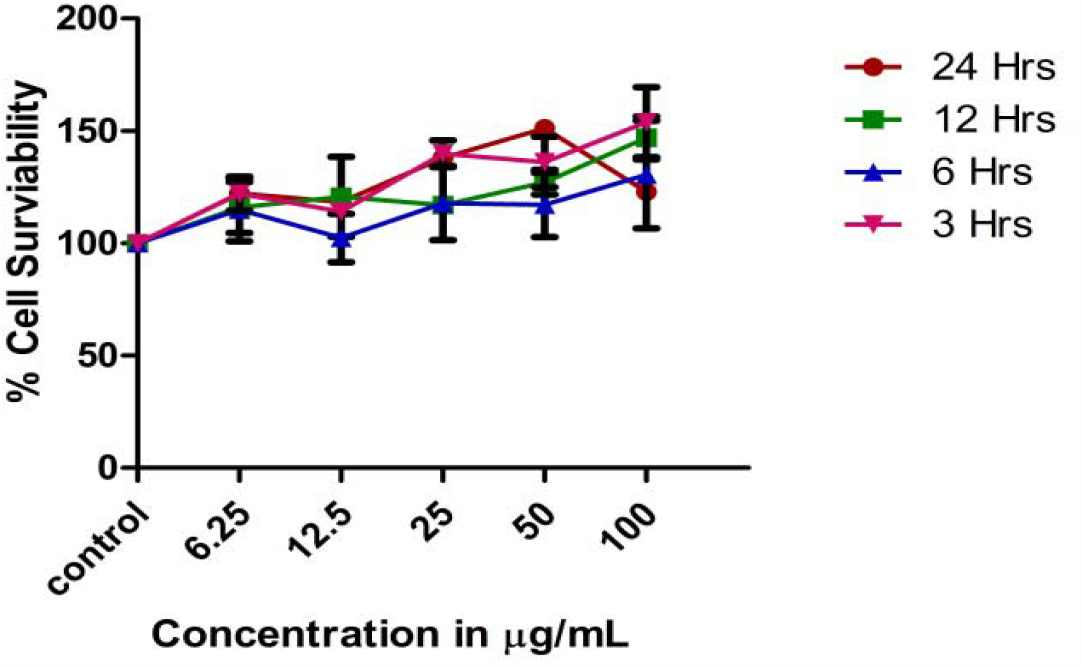
Cell Proliferation assay on RAW 264.7 cell lines at different time points

Various chemicals and metabolites established their role in inhibiting tumour growth and its metastasis. Likewise, many lectins were reported as anti-tumour agents in treating different cancers. They are playing an important role in inhibiting tumour growth and metastasis by involving or influencing the autophagy and apoptosis [31].

Eutirucallin, a latex lectin from *Euphorbia tirucalli* exhibited potential anti-tumour activity against various cancer cell lines [1]. ECL also exhibited anti-tumour activity against HeLa cell lines and inhibited cell migration (Fig. 6). Cell proliferation was observed with non-tumour cells (J774A.1) whereas cell death was noticed in tumour cells (HeLa). Anti-tumour activity was observed more as increasing the concentration where the opposite effect was shown with non-tumour cells. The ability of cell migration was also ceased by lectin with all concentrations and exhibited maximum anti-tumour activity at 200 μg/ml. These results promise the anti-tumour activity of the lectin from *E. caducifolia* latex.

**Fig. 6.**
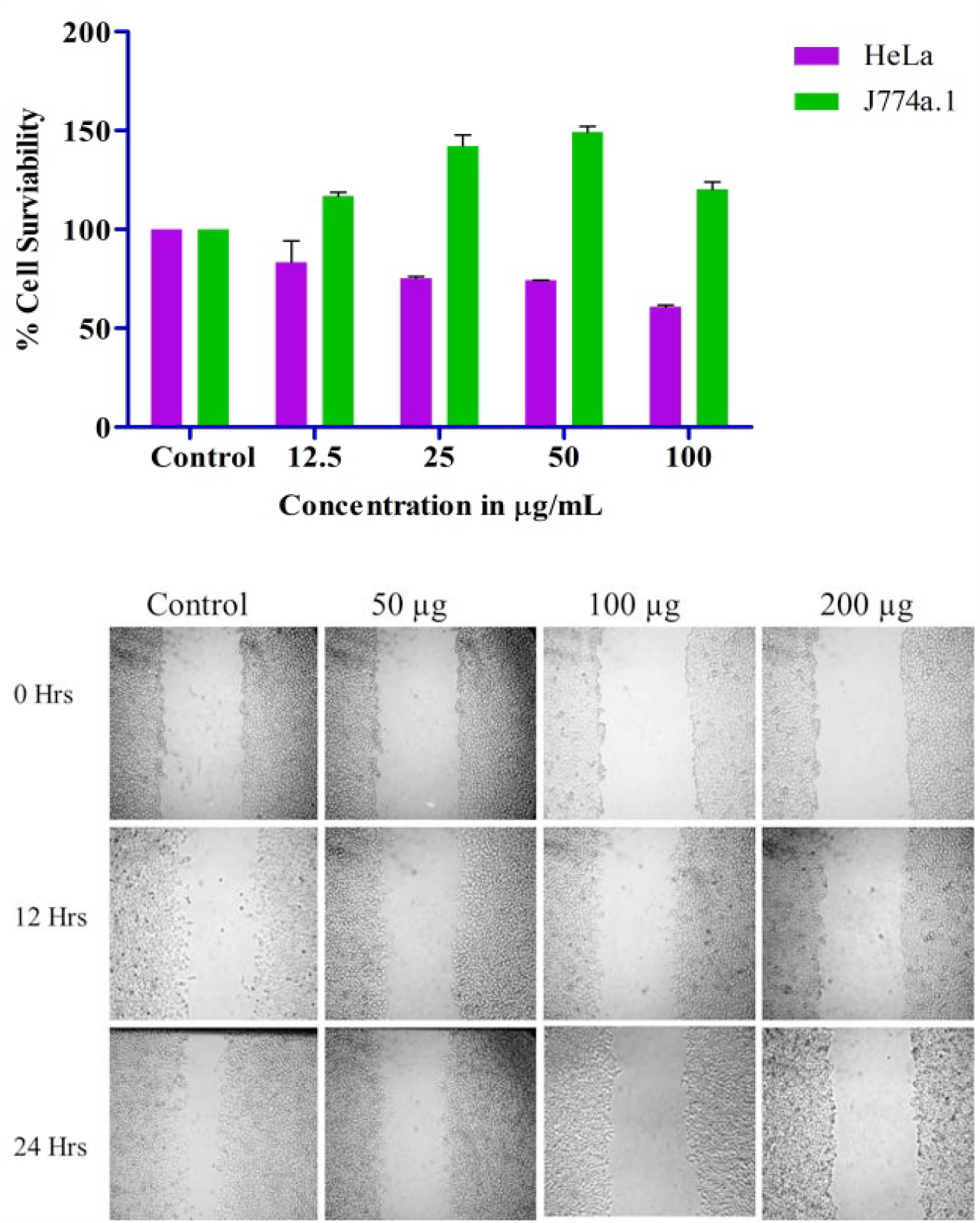
Anti-tumour activity on Hela cells after 24 h of treatment and Inhibition of HeLa cells migration

## 4. Conclusion

Our results suggest that *E. caducifolia* latex contains galactose specific lectin with mitogenic nature on human macrophages (RAW 264.7) and anti-tumor activity on cancer cells (HeLa). Lectin is predominant with β conformation structure and active at pH 6-8 and 60-70°C. Moreover, CLLSG matrix yields more purified lectin than classical CLGG.

## Acknowledgements

Kusuma Venumadhav acknowledges to Rajiv Gandhi National Fellowship, University Grant Commission (UGC), New Delhi, India for providing financial assistance as a Senior Research Fellow.

## Conflict of interest

The authors have declared no conflict of interest.

## Notes

### Competing Interest Statement

The authors have declared no competing interest.

### Summary of Updates

Manuscript hasbeen revised with certain relevant figures

## References

[1] Santiago, F.M., Palharini, J.G., Richter, A.C., Silva, M.F., et al., Eutirucallin: A Lectin with Antitumor and Antimicrobial Properties. Front. Cell. Infect. Microbiol. 2017, 7, 1–13.

[2] Stillmark, Uber rizin, ein giftiges ferment aus dem samen von Ricinus communis L. und einigen anderen Euphorbiaxeen. Ina. Dorpat 1888.

[3] Mellor, R.B., Gadd, G.M., Rowell, P., Stewart, W.D.P., A phytohaemagglutinin from the Azolla-anabaena symbiosis. Biochem. Biophys. Res. Commun. 1981, 99, 1348–1353.

[4] Suzuki, I., Satto, H., Inoue, S., Migita, S., Takahashi, T., Purification and Characterization of Two Lectins from Aloe arborescens Mill. Biochem 1979, 85, 163–171.

[5] W G Carter, M E Etzler, Isolation, characterization, and subunit structures of multiple forms of Dolichos biflorus lectin. J. Biol. Chem. 1975, 250, 2756–2762.

[6] Lamb, J.E., Shibata, S., Goldstein, I.J., Purification and Characterization of Griffonia simplicifolia Leaf Lectins. Plant Physiol. 1983, 71, 879–87.

[7] Horejsi, V., Haskovec, C., Kocourek, J., Studies on lectins XXXVIII: Isolation and characterization of lectin from locust bak (Robinia pseudacacia L.). Biochim. Biophys. Acta 1978, 532, 98–104.

[8] Shibuyas, N., Goldstein$, I.J., Broekaertg, W.F., Nsimba-Lubakiq, M., et al., The Elderberry (Sambucus nigra L.) Bark Lectin Recognizes the Neu5Ac(a2-6)Gal/GalNAc Sequence*. J. Biol. Chem. 1987, 262, 1596–1601.

[9] Gietl, C., Kauss, H., Ziegler, H., Affinity Chromatography of a Lectin from Robinia pseudoacacia L. and Demonstration of Lectins in Sieve-Tube Sap from Other Tree Species. Planta 1979, 144, 367–371.

[10] Van Damme, E.J.M., Allen, A.K., Peumans, W.J., Isolation and characterization of a lectin with exclusive specificity towards mannose from snowdrop (Galanthus nivalis) bulbs. FEBS Lett. 1987, 215, 140–144.

[11] Damme, E.J.M. Van, Allen, A.K., Peumans, W.J., Related mannose-specific lectins from different species of the family Amaryllidaceae. Physiol. Plant. 1988, 73, 52–57.

[12] Damme, E.J.M., Peumans, W.J., Isolectins in Narcissus: complexity, inter- and intraspecies differences and developmental control. Physiol. Plant. 1990, 79, 1–6.

[13] Oda, Y., Minami, K., Isolation and characterization of a lectin from tulip bulbs, Tulipa gesneriana. Eur. J. Biochem. 1986, 159, 239–245.

[14] Peumans, W.J., Nsimba-Lubaki, M., Carlier, A.R., Van Driessche, E., A lectin from Bryonia dioica root stocks. Planta 1984, 160, 222–228.

[15] Yeung, H.W., Wong, D.M., Li, W.W., in:, 4 th Asian symp. Med. plants species, Mahidol. Unive, Bankok, Thailand 1980, p. 94.

[16] Venumadhav, K., Seshagirirao, K., Phytochemical screening and Antioxidant activity of Euphorbia caducifolia extracts n.d.

[17] Seshagirirao, K., Prasad, M.N.V., in:, Saxena P. (Ed.), Dev. Plant-Based Med. Conserv. Effic. Saftey, Kluwer Academic Publishers, Netherlands 2001, pp. 199–210.

[18] Seshagirirao, K., Leelavathi, C., Sasidhar, V., Cross-linked leucaena seed gum matrix: an affinity chromatography tool for galactose-specific lectins. J. Biochem. Mol. Biol. 2005, 38, 370–2.

[19] DuBois, M., Gilles, K. a., Hamilton, J.K., Rebers, P. a., Smith, F., Colorimetric method for determination of sugars and related substances. Anal. Chem. 1956, 28, 350–356.

[20] Pitthard, V., Finch, P., GC-MS Analysis of Monosaccharide Mixtures as their Diethyldithioacetal Derivatives[: Application to Plant Gums Used in Art Works. Chromatogr. Suppl. 2001, 53, 317–321.

[21] Lis, H., Sharon, N., Soy bean (Glycine max) agglutinin. Method. Enzym. 1972, 28B, 360–365.

[22] Khan, N., Rahim, S.S., Boddupalli, C.S., Ghousunnissa, S., et al., Hydrogen peroxide inhibits IL-12 p40 induction in macrophages by inhibiting c-rel translocation to the nucleus through activation of calmodulin protein. Blood 2006, 107, 1513–1520.

[23] Lynn, K.R., Clevette-Radford, N.A., Lectins from latices of Euphorbia and Elaeophorbia species. Phytochemistry 1986, 25, 1553–1557.

[24] Barbieri, L., Falasca, A., Franceschi, C., Licastro, F., et al., Purification and properties of two lectins from the latex of the euphorbiaceous plants Hura crepitans L. (sand-box tree) and Euphorbia characias L. (Mediterranean spurge). Biochem. J. 1983, 215, 433–9.

[25] Lynn, K.R., Clevette-Radford, N.A., Isolation and characterization of proteases from Euphorbia lactea and Euphorbia lactea cristata. Phytochemistry 1986, 25, 807–810.

[26] Duk, M., Lisowska, E., Effect of pH on the binding of Ecia graminea lectin to erythrocytes Dependence on the chemical character of red-cell receptors 1984, 78, 73–78.

[27] Fosgerau, K., Hoffmann, T., Peptide therapeutics: Current status and future directions. Drug Discov. Today 2015, 20, 122–128.

[28] Ashraf, M.T., Khan, R.H., Mitogenic lectins. Med. Sci. Monit. 2003, 9, RA265–9.

[29] Stirpe, F., Licastro, F., Morini, M.C., Parente, A., et al., Purification and partial characterization of a mitogenic lectin from the latex of Euphorbia marginata. BBA - Gen. Subj. 1993, 1158, 33–39.

[30] Kitajima, S., Miura, K., Aoki, W., Yamato, K.T., et al., Transcriptome and proteome analyses provide insight into laticifer’s defense of Euphorbia tirucalli against pests. Plant Physiol. Biochem. 2016, 108, 434–446.

[31] Yau, T., Dan, X., Ng, C.C.W., Ng, T.B., Lectins with potential for anti-cancer therapy. Molecules 2015, 20, 3791–3810.

